# Neural dynamics of accumulating and updating linguistic knowledge structures

**DOI:** 10.1101/495168

**Authors:** Ruud M.W.J. Berkers, Marieke van der Linden, David A. Neville, Marlieke T.R. van Kesteren, Richard G.M. Morris, Jaap M.J. Murre, Guillén Fernández

## Abstract

Knowledge is acquired by generalization and integration across learning experiences, which can then be applied to future instances. This study provides novel insights into how linguistic associative knowledge is acquired by systematically tracking schematic knowledge formation while participants were learning an abstract artificial language organized by higher-order associative regularity. During learning, we found activity in the left inferior frontal gyrus in response to knowledge updating during feedback presentation, as well as in response to available accumulated knowledge during retrieval. A complementary signal was found in the caudate nucleus, where activity correlated with the availability of recently acquired knowledge during retrieval, suggesting it initially supports the retrieval of knowledge. Furthermore, we find that activity in a set of regions, including the medial prefrontal cortex and hippocampus, scaled with accumulated knowledge during feedback presentation, which might be indicative of increased generalization of features of the hierarchical knowledge structure. Together, these results provide a mechanistic insight into how linguistic associative knowledge is acquired by generalization across repeated learning experiences.

## Introduction

Associative knowledge structures, or schemas, capture consistent relationships amongst low-level perceptual features as well as higher-order concepts across multiple episodes (van Kesteren et al. 2012; Ghosh and Gilboa 2014). Humans are driven towards discovering structure across seemingly arbitrary low-level contingencies. In one demonstration, people studied geometric figures linked to arbitrary letter strings (Kirby et al. 2008). When cued with the geometric figures and asked to type the associated label, they were generally poorly reproduced. Interestingly, when using one participants’ output as labels for the next participant, and repeating this process iteratively, an artificial language evolved with a higher-order hierarchical structure. This schematic structure was imposed by iterative errors that drove the language to attain compositionality. One example of such an evolved language involved a structure whereby each syllable uniquely denoted a perceptual feature: the first syllable denoting the colour; the second, geometric shape; and the last syllable denoting the movement trajectory of the figure (see left panel of figure 1). Moreover, the language developed a structure that was consistent across individual exemplars whilst remaining uniquely identifiable for individual exemplars.

**Figure 1.**
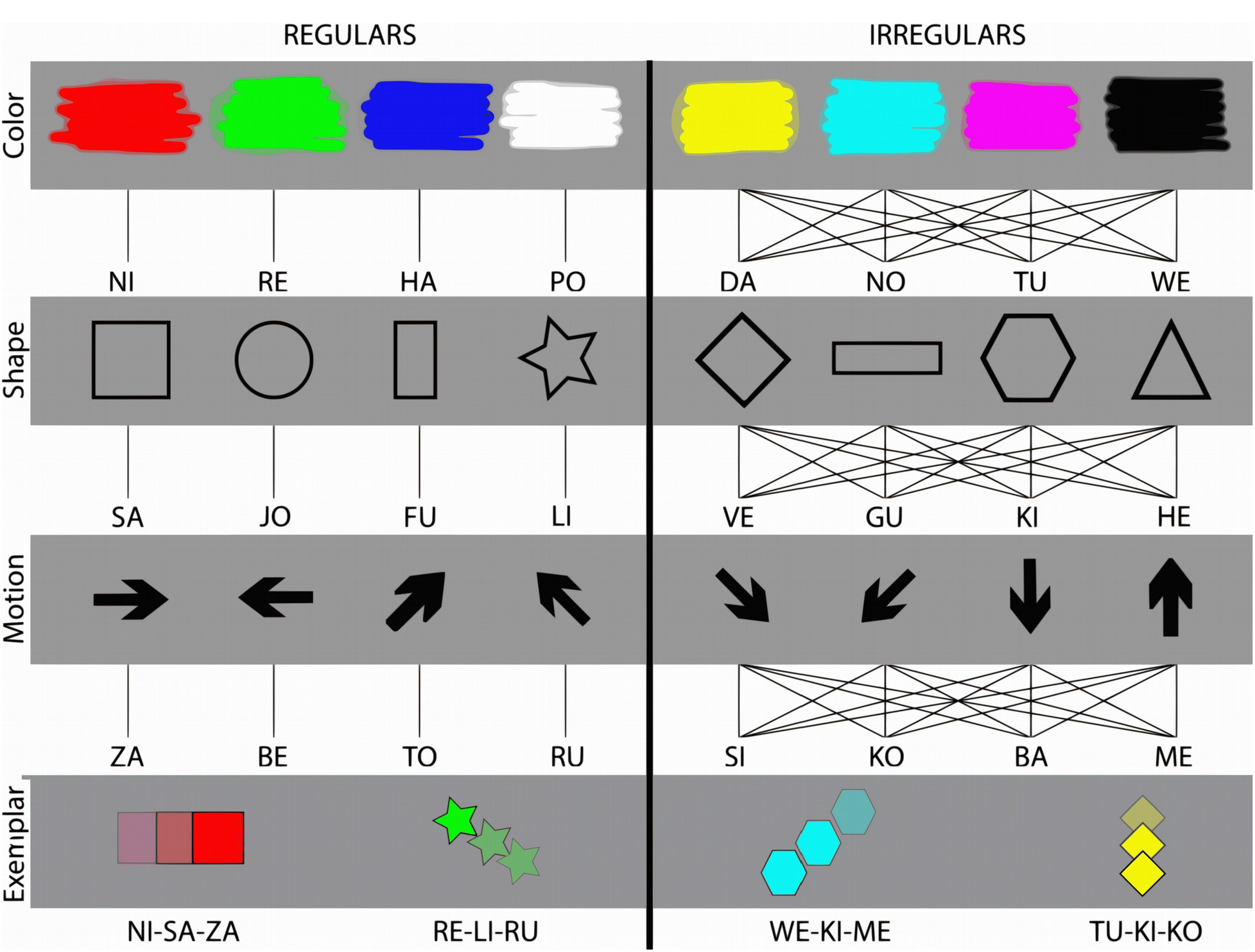
Stimulus materials. Subjects were presented with pairs of geometric figures and artificial word labels. The figures consisted of a defining colour, shape and movement, and the words consisted of three corresponding syllables that uniquely mapped onto the objects’ perceptual features. In the regular set, a particular instantiation of a perceptual feature was always associated with the same syllable (left panel), whereas in the irregular set the same feature instantiation could be associated with different syllables, depending on the exemplar configuration (right panel). Below: four examples of associations between figures and artificial words from the regular set (left panel) and the irregular set (right panel) are shown.

Schematic knowledge is thought to be stored in associative neocortical structures (Bartlett 1932; van Kesteren et al. 2012; Ghosh and Gilboa 2014; Wagner et al. 2015; van der Linden et al. 2017), consisting of categorical and hierarchical nodes (Miller et al. 2002; Barsalou 2009). Schematic knowledge is acquired by extracting regularities across episodes to build a structure of higher-order relationships that can then be applied to novel instances (Brady and Oliva 2008). Often, activated prior knowledge constrains the acquisition of novel, related knowledge (Markman and Hutchinson 1984), and this learning benefit is associated with hippocampal and medial prefrontal processing (Tse et al. 2007, 2011, van Kesteren et al. 2010, 2014). Without prior constraints, however, humans typically require many trials to acquire generalizable knowledge structures (Seger et al. 2000; Seger and Cincotta 2006). In contrast with the hippocampal memory system that stores event-specific information, the neocortex accumulates generalizable knowledge across multiple similar events to build the general statistical structure of environmental relations (McClelland et al. 1995; O’Reilly and Norman 2002). A similar relationship is found in corticostriatal loops during feedback-based learning of stimulus-response associations, with the basal ganglia responding on a trial-by-trial basis to reward-prediction-errors (reward-gated plasticity) and the neocortex responding to the accumulation of reward across repetitions (reward-shaded plasticity; Seger and Miller 2010). Both the striatum and hippocampus thus learn on the basis of unique exposures to exemplars (Daw et al. 2005; Seger and Cincotta 2006) in contrast with a slower learning neocortex (Pasupathy and Miller 2005). The distinct contribution of the striatum and the hippocampus is often subtle as demonstrated by research on the probabilistic classification task. Here, patient studies suggest learning to be dependent on the striatum (Holl et al. 2012; Dalton et al. 2013) and the hippocampus (Knowlton et al. 1994). Imaging studies report either a trade-off with initial hippocampal activity and a slower build-up of prolonged activity in the caudate nucleus of the striatum (Poldrack et al. 2001), or both striatal and hippocampal activity tracking initial learning (Kumaran et al. 2009). In the latter study, the pattern across many single associations contained a higher-order structure consisting of two associative rules, and generalization of this structure was related to initial learning-related activity in the hippocampus and connectivity between hippocampus and medial prefrontal cortex (Kumaran et al. 2009). Other imaging studies of initial knowledge acquisition have also typically used tasks with one or two associative rules and dichotomous choice options (Seger et al. 2000; Seger and Cincotta 2006). However, from these studies it is not clear how activity dynamically unfolds across learning in the striatum, hippocampus and neocortical knowledge representation areas.

This fMRI study uses a novel learning paradigm consisting of a linguistic hierarchical knowledge structure that is gradually acquired (inspired by Kirby et al. 2008), allowing us to model trial-by-trial knowledge build-up across a number of interleaved trials within a single learning session. Critically, participants learned associations between geometric figures on the one hand (defined by their colour, shape and movement) and tri-syllabic word strings on the other hand. These associations contained higher-order regularities in their mappings across exemplars. Exemplar associations were replaced with new exemplars halfway through the learning session, allowing the establishment of generalization across trials (see figure 1). We then tracked learning from an initial state, where associations are deemed arbitrary, to an end-state where these associations have acquired meaning within the higher-order associative structure. The State-Space model (Smith et al. 2004) was then used to systematically track accumulation and updating of knowledge during the acquisition session during the cue presentation (test phase) and feedback presentation (learning phase). This approach allows us to closely track how various brain regions dynamically contribute to the gradual acquisition of a complex linguistic knowledge structure.

## Materials and Methods

### Participants

Thirty-two healthy, right-handed subjects with normal or corrected-to-normal vision participated in the experiment (age range: 19-32 years; 20 female). Subjects received monetary compensation for participation and could earn extra money based on performance. One subject was excluded from further analysis due to scanner malfunction, and five subjects were excluded because they failed to reach the learning criterion defined by reaching the critical learning trial (CLT, see Behavioural Analysis) over the course of the experiment. One further participant was excluded from the MRI-analysis due to excessive movement inside the scanner. All subjects provided written informed consent. The study was conducted according to a protocol approved by the local review board (CMO Region Arnhem-Nijmegen, the Netherlands).

### Stimuli

The stimuli consisted of geometric figures described by artificial tri-syllabic word labels. The figures and word labels did not have a meaningful association to real-life figures and words. As such, the influence of prior knowledge on learning the associations was minimized. More specifically, the stimuli consisted of visual geometric figures defined by a specific colour, shape, and movement trajectory across the screen. There were eight isoluminant colours (red, green, blue, white, yellow, cyan, magenta, and black) presented on an isoluminant grey background, eight different shapes (square, circular, vertical rectangle, star, diamonds, vertical rectangle, hexagon, and triangle, all made to fit into a rectangle of 142 pixels), and eight movement trajectories (all moving at two pixels per screen refresh along a polar angle of respectively 0, 45, 90, 135, 180, 225, 270 and 315 degrees). In total, these three features could be combined into 512 unique exemplars. Similarly, 512 unique word labels could be made on the basis of the 24 syllables used (see figure 1).

Two sets of associations were used for the acquisition session. Both sets contained four colours, shapes, and movements and each particular perceptual feature was only used in one of the two sets. Therefore, each set contained 64 exemplars with a unique configuration of three perceptual features (colour, shape and movement). The syllables were distributed across both sets, each set containing 64 word exemplars with a unique configuration of three syllables. The regular set contained associative regularities across exemplars that could be discovered and generalized to novel exemplars. Each visual feature was consistently associated with a syllable in a fixed position within the tri-syllabic word (colour corresponded to the first, shape to the second, and movement to the third syllable). Furthermore, each instantiation of a feature corresponded to an uppercase syllable (e.g. red = ‘NI’, square geometry = ‘SA’, rightward movement = ‘ZA’, fig. 1). In contrast, the irregular set contained no regularity across individual exemplars. Here, in different exemplars, features (colours, shapes and movements) were paired with different syllable locations, and feature instantiations (e.g. ‘red’, ‘square’ and ‘rightward movement’) were paired with different syllables. Here, the pairings between figure and word labels were consistent only within repetitions of the same exemplar, but not in different exemplars with overlapping features (see fig. 1 for examples). Colours, geometric shapes, movements and syllables were assigned to the two sets in a counterbalanced manner across participants.

### Task and procedures

Participants were instructed that they were to learn a new language consisting of geometric figures denoted by a tri-syllabic word label. They were informed that there was regularity in the associations between figures and artificial words, but the nature of this regularity was not disclosed. Participants viewed blocks consisting of either geometric figures or word labels, such that all perceptual features and syllables had been seen across this pre-exposure period. This served to familiarize participants with the stimuli on a perceptual level and to rule out stimulus novelty effects during the ensuing scan session. To become familiarized with the requirements of the task, participants were briefly trained on nine trials similar to those that were presented later during the acquisition session. This training used colours, shapes, movements and syllables that were randomly sampled from the entire set.

During the scanned acquisition session participants were presented with trials from both regular and irregular sets distributed across 8 blocks (see figure 2). During the first four blocks, twelve regular figure-word combinations and four irregular figure-word combinations were repeated in each block. In the second set of four blocks, 12 new regular and four new irregular combinations were introduced and repeated across the remaining blocks. As such, participants’ ability to generalize across exemplars of block 4 and 5 could be assessed. The order of regular and irregular trials within a block was randomized. In total, 96 trials from the regular set and 32 trials from the irregular set were presented across four runs (each run contained two blocks). In each trial, the figure was presented for 3s (the test phase), then participants were asked to select the corresponding tri-syllabic word from three options per syllable (the response phase, duration 6s), and lastly the correct response was presented (feedback phase, duration 3s). Participants were instructed to retrieve the word label immediately upon seeing the figure and to maintain that information until they could give the correct response (see figure 2).

**Figure 2.**
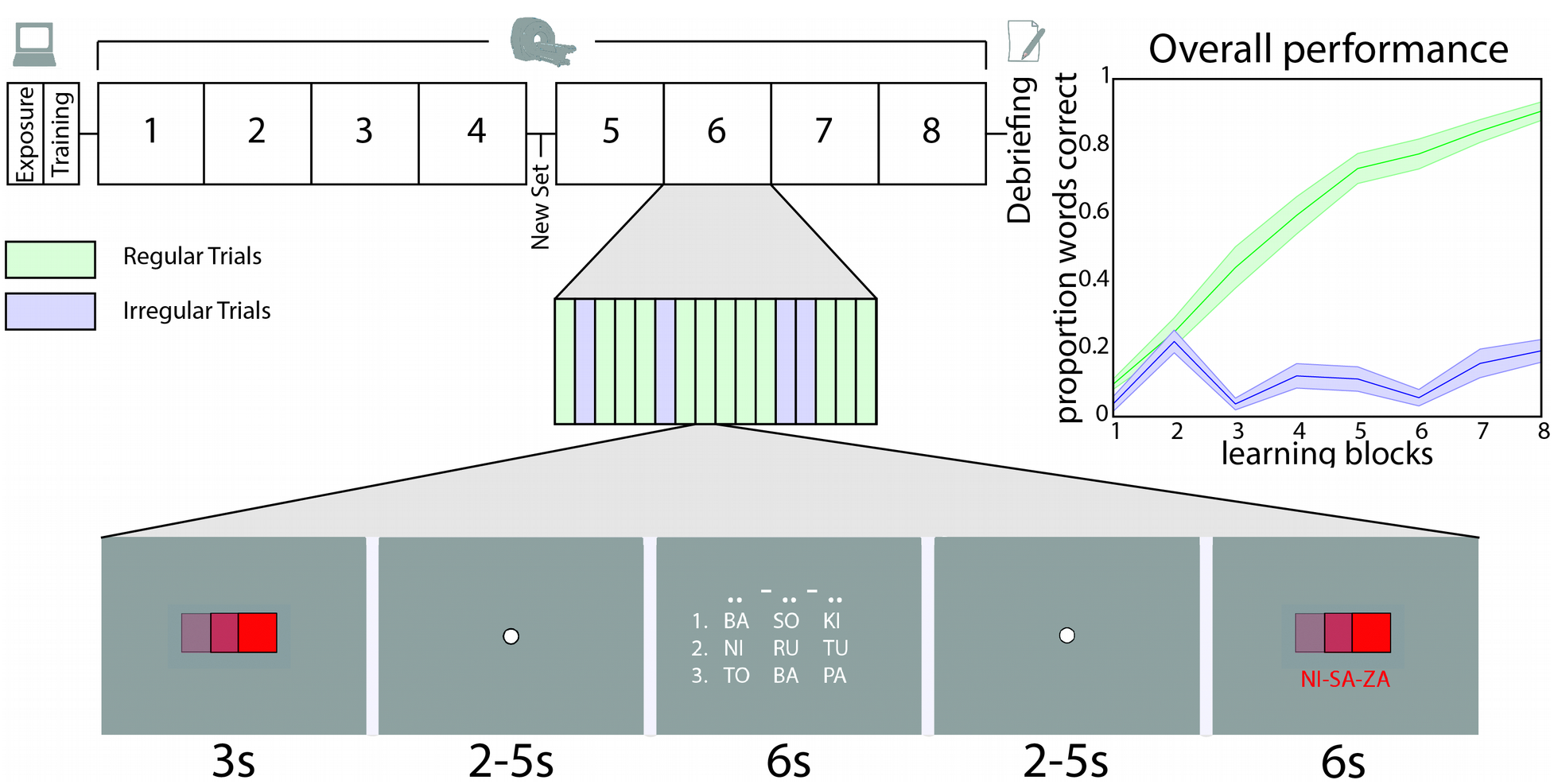
Task design and task performance. After pre-exposure to stimulus materials and pre-training, participants learned figure-word associations across eight blocks. Each block contained 12 regular trials and 4 irregular trials, these were repeated four times. After the first four blocks, the exemplars were switched. In each trial, first the figure was presented (the test phase, duration 3s), then the corresponding tri-syllabic word was selected (the response phase, duration 6s), and lastly the correct response was presented (learning phase, duration 3s). Participants were instructed to retrieve the word label upon seeing the cue, and maintain that information until the correct response could be given. Inset: Plot of overall performance for irregular and regular trials demonstrates a steadily improving learning curve for regular trials.

During the response-phase, participants were asked to select three syllables comprising the complete word, by selecting one out of three alternatives for each syllable location (first, second, and third syllable of the word). The chance level for having one individual syllable correct is therefore 1/3, and the chance level for having the entire word correct is 1/27 (1/3 * 1/3 * 1/3). A blue bar underlined the syllable location where at each moment a syllable needed to be selected (first, second, and third syllable of the word). To reduce interference across sets, the irregular set was clearly distinguishable from the regular set. Specifically, the three syllable alternatives were underlined with light-blue bars, such that participants were able to distinguish the regular and irregular sets. The regular and irregular trials were similar, but participants could not generalize learned associations across exemplars. For the alternative response options, syllables were pseudorandomly selected from all syllables that occurred across regular and irregular associations at that particular syllable location (first, second, and third syllable of the word). This ensured that both syllables from the regular and irregular set were repeated to a roughly equal extent across learning, and effects of syllable familiarity were ruled out.

In the feedback-phase, the figure was presented again with the correct word underneath, presented in red if one of the selected syllables had been incorrect and green if the entire word was correct. Participants were instructed to compare feedback with their initial response and learn from the discrepancies between response and feedback. All trials and phases within a trial were separated by a jittered interval of 2-5 seconds. During the entire experiment the participants were instructed to fixate on a white fixation dot that remained visible at the centre of the screen. After each block, a baseline block with a jittered length of 9-11 seconds was presented. The stimuli were presented using a projector at the rear of the scanner bore (60 Hz refresh rate, 1024 by 768 resolution) viewed by the participants through a mirror attached to the headcoil (covering 6° of horizontal and 7° of vertical visual angle). The experiment was programmed in Matlab, using the Psychophysics Toolbox extensions (Brainard 1997).

Following completion of the acquisition session, participants were debriefed to assess whether they had discovered the organizing principles underlying the associations. They were first asked to provide the meaning of all 24 syllables from the regular set (e.g. by writing ‘red’ for colour, writing ‘hexagon’ or making a small drawing for shape, and ‘to the right’ or an arrow for movement). Next, they were asked whether they knew which perceptual feature corresponded to the first, second, and third syllable in the word label. As such, this questionnaire probed explicit knowledge of the associative regularity inherent in the regular set.

### Behavioural analysis

In the current paradigm, each word label consisted of a sequence of three syllables. The participant was presented with three response options per syllable (33 % chance probability of selecting a correct syllable) and thus with a total of 27 possible word responses per trial (~4% chance probability of correct response for the entire word). Vectors coding (0 to 3 for the number of features correct) for trial-by-trial performance across all 128 trials were extracted for individual subjects.

To track the state of knowledge across the acquisition session, individual learning curves were estimated using the State-Space model (Smith et al. 2004). In this model the accumulation of knowledge is described as an increase in the probability of a correct response across trials. The model is characterized by two equations: an observation equation and a state equation. The observation equation describes how the observed binary choice data (e.g. ‘correct’ or ‘incorrect’) relates to a hidden state or latent learning process. Each trial involves a sequence of three responses to select the correct syllable for each perceptual feature. Therefore, the observation equation is best characterized as a binomial process (i.e. a sequence of three Bernoulli trials). The state equation describes the hidden learning process that evolves across trials and is defined as a Gaussian random-walk process. Therefore, learning in this model is reflected by an increase in the latent state process that ultimately leads to an increase in the number of correct responses (i.e. higher probability of selecting the correct syllable for each feature). In the state-space model, inferences on the learning process are made from the perspective of an ideal observer, using the complete sequence of trials to estimate the time-course of learning. The state-space model was fitted to the data using numerical optimization techniques based on the Expectation Maximization (EM) algorithm (code obtained from www.neurostat.mit.edu).

To verify that the state-space model provided the best account of the behavioural data, three alternative models were also fitted to the learning data: the Rescorla-Wagner model (Rescorla and Wagner 1972), the learning component of the memory chain model (MCM; Chessa and Murre 2007; Murre 2013) and the moving average model (MA; Smith et al. 2004). The models were fitted to the data using customized code adapted from freely available online resources and Maximum Likelihood fitting routines. Fitting of the state-space model was carried out using custom code for MATLAB^®^ (adapted from Smith et al., 2004). Fitting of the Rescorla-Wagner model was done by calculating the equilibria of the model (Danks 2003) using functions implemented in the R^®^ package ‘ndl’ (Arppe et al. 2015). Fitting of the MCM model was carried out using custom code and standard optimization routines for Mathematica®. Last, the moving average model was fitted to the data using the toolbox Forecast (Hyndman et al. 2013) for R^®^.

Model selection was carried out by calculating Bayesian Information Criterion scores (Schwarz 1978) for each model, which is a measure of the goodness-of-fit that penalizes for the number of free parameters. Therefore, the model with the lowest score represents the most parsimonious account of the data. In line with previous studies (Kumaran et al. 2009), results indicated that the state-space model provided the best description of the observed data (see table 1). The fit of learning curves generated by the moving average method was worse than the state-space model, but superior to the Rescorla-Wagner model. The memory chain model performed better than the RW, but worse than the state-space model or moving-average model. The state-space model thus provides the most accurate description of the experimentally observed individual learning curves and therefore the estimated ‘p mode’ parameter (the mode of the distribution of estimated probabilities per trial) from the state-space-model was used as model parameter for parametric model-based fMRI analyses. Furthermore, the critical learning trial (CLT, see Smith et al. 2004) was defined as the first trial where it can be concluded with reasonable certainty that a subject performs better than a certain threshold (95% confidence interval exceeds chance performance). This threshold performance was here defined as 66%, which corresponds roughly to having two syllables correct.

**Table 1.**
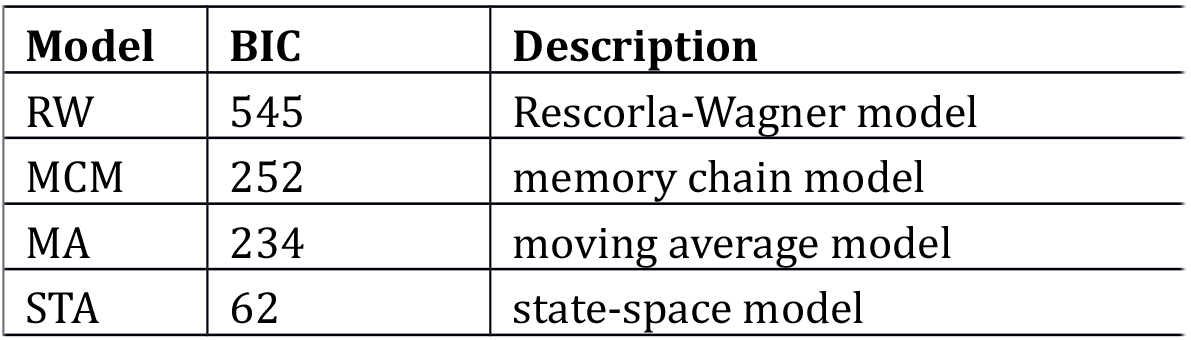
Model comparison. Bayesian Information Criterion (BIC) scores for each of the four learning models. The model with the lowest score represents the most parsimonious account of the data.

### Imaging parameters and acquisition

T2*-weighted echo planar images (EPI) with BOLD (blood-oxygen-level-dependent) contrast were acquired on a 3 Tesla Siemens Skyra MRI scanner. We scanned 45 oblique axial slices angled at 30° in the anterior-posterior axis, TR 2.44 s, 2 mm thickness (0.5 mm gap), in-plane resolution 2.5 × 2.5 mm, field-of-view 212 mm. To minimize signal dropout in the temporal lobes and medial prefrontal cortex a dual-echo sequence (Halai et al. 2015) was used (TE1 = 15 ms and TE2 = 36). Additionally, a structural T1-weighted 3D magnetization prepared rapid acquisition (MPRAGE) gradient echo sequence image (192 slices, voxel size =1 × 1 × 1 mm) was acquired for each participant.

### fMRI data preprocessing

The first six volumes were discarded to permit T1 relaxation, and the next thirty volumes were used to calculate the weighting for recombining the two echo images. The images within a run were realigned to the first image, and subsequently the two echo images for each volume were recombined using custom scripts and weighting parameters (Poser et al. 2006). Subsequently, images were preprocessed using the statistical parametric mapping software SPM (http://www.fil.ion.ucl.ac.uk/spm/) and custom scripts written in Matlab (http://www.mathworks.com/products/matlab). Next, structural and functional images were segmented and normalized to MNI-space on the basis of their grey and white matter templates using DARTEL. Compartment maps were generated for grey matter, white matter and CSF, and nonspecific nuisance regressors were created modelling signal from white matter and CSF-compartments. Normalized images were smoothed using a Gaussian kernel with full-width half-maximum of 5 mm. Custom scripts were used to detect and remove spike artifacts.

### FMRI analyses

The fMRI data was analyzed in SPM12 using the general linear model (GLM), estimated in two stages. Subject-specific experimental effects were modelled at the first-level using GLMs. Then, a random-effects analysis was performed at the second level using one-sample t-tests, resulting in group-level statistical parametric maps. Task regressors were included in GLMs to model the cue-phase, response-phase and feedback-phase separately for regular and irregular trials, while the inter-trial interval and baseline blocks served as implicit baseline. Events were modelled as a boxcar function and convolved with the canonical haemodynamic response function (HRF). Furthermore, button presses were modelled as stick functions and convolved with the canonical haemodynamic response function (HRF). Movement parameters were included as regressors of no interest. To deal with non-specific signal fluctuations, signal time-courses were extracted from white-matter and cerebrospinal fluid compartments and included as regressors of no interest. We employed a high-pass filter with a low cut-off of 1/835 Hz, as the task power spectrum revealed that a more conventional cut-off of 1/128 Hz removed gradual learning-related fluctuations in the BOLD-signal. Runs were also concatenated to prevent the removal of gradual learning signals. Temporal autocorrelation was modelled using AR(1).

Analyses of regional activation focused on parametric modulations, namely regressors in GLMs to detect brain regions where BOLD-activity was modulated by the trial-by-trial state of knowledge during regular trials (the irregular trials were not considered here as the amount of trials was low). The first model included subject-specific vectors denoting the probability of a correct response on any given trial (probability range: 0-1) as estimated by a state-space model (Smith et al. 2004). This vector is a proxy for the level of knowledge accumulated at any given trial, and thus approximates the amount of knowledge that could be retrieved at any trial (see figure 3a). The accumulation function was included as a parametric modulator in the GLM to model the cue-period, as a higher value here approximates the amount of knowledge that can be retrieved. During the feedback-period, knowledge is not actively retrieved, and once more knowledge has accumulated less updating occurs. Therefore, activity related to accumulated knowledge during feedback presentation might represent a variety of factors (see Discussion). The second model included the difference function of the first regressor (the state-space learning curve), which indicates the change in probability of a correct response on each trial compared to the previous trial (see figure 3b). The updating parameter was used to model the cue period, as it is an estimate for recently acquired knowledge that can be recruited in the current test phase. This function was given the value of zero for the first trial, where no knowledge was present by definition. The same updating function was used to model the feedback period, but shifted by one trial earlier in the sequence (such that the updating function represents the change in knowledge on the current trial compared to the next), and the updating function was appended with a 0 in the last trial. Both model 1 and model 2 thus included 10 task regressors (test-phase, response-phase and feedback-phase for regular and irregular trials, button presses, parametric modulator for test-phase, response-phase and feedback-phase of regular trials), along with the compartment and movement regressors. Condition-specific experimental effects (regression coefficients) were obtained in a voxel-wise manner for each participant. At the second (random-effects) level, participant-specific linear contrasts of the parameter estimates were entered in a one-sample t-test to construct the group-level statistical map. We considered results that survived correction for multiple comparisons (family-wise error (FWE) correction) at p<0.05 across the whole brain or within independently defined anatomical masks (small-volume correction, SVC) after initial thresholding at p=0.001 uncorrected. Anatomical masks were defined using the WFU PickAtlas Tool 2.4 as implemented in SPM.

**Figure 3.**
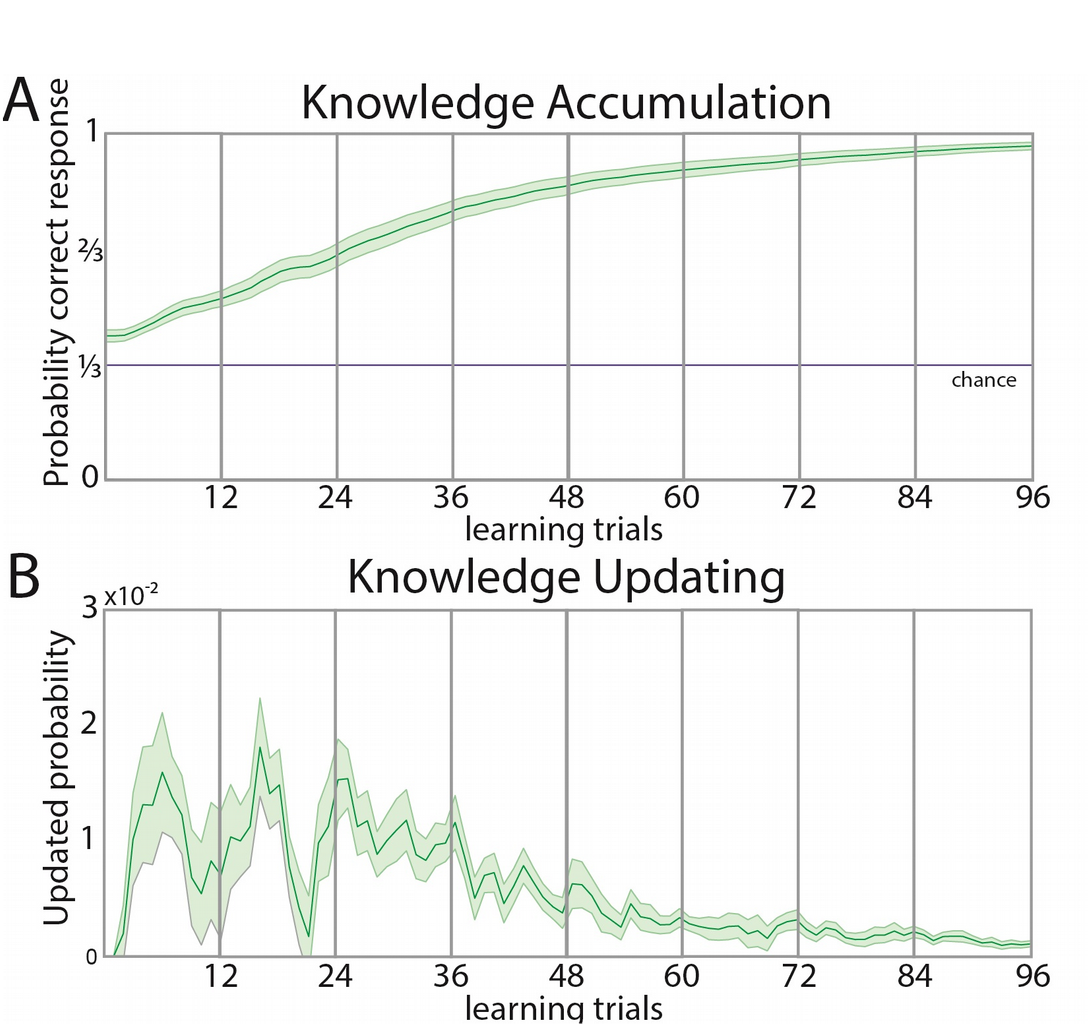
Learning parameters estimated from the State-Space model. Top panel: Group-averaged probability of making a correct response on each trial as estimated by the State-Space model. Bottom panel: Averaged change of the probability of making a correct response on each trial compared to the previous trial, as estimated by the State-Space model.

### Beta-series extraction

To characterize the temporal dynamics of regional activity across learning, beta-values were analysed from regions of interest across all learning trials. Specifically, trial-specific beta-images were obtained for all cue- and feedback-presentations by running one GLM with separate columns for each cue-presentation. A second GLM was run with separate columns for each feedback-presentation. The resulting beta-images were sorted according to their temporal trial order for cue and feedback presentations, and signal was extracted across beta-images for each ROI. Furthermore, to control for subject-specific differences in the learning rate, the timeseries were also centered around the critical learning trial. These subject-specific timeseries were then smoothed using a centered moving average with a sliding window of three timepoints, and the resulting vectors were then averaged across all participants.

## Results

### Behavioural learning

Participants studied object-word associations across 128 trials in eight scanner blocks of 16 trials. Each block contained 16 unique associations that were repeated across the first four blocks, after which the associations were changed and a different set of 16 associations was repeated across the next four blocks. Twenty-six participants reached the critical learning trial (mean CTL = 39.96, SD = 19.12), and exceeded chance level performance on the last block (proportion of all words correct on last block, mean = 0.74, SD = 0.11, chance level = 0.037, t_(25)_ = 33.54, *p* < 0.001). Each block contained twelve exemplars from the regular set and four exemplars from the irregular set. In both conditions, learning was evident by above-chance performance in the eighth and last block (proportion regular correct, mean = 0.92, SD = 0.12, versus chance level = 0.037, *t*_(25)_ = 37.84, *p* < 0.001, mean proportion irregular correct, mean = 0.19, SD = 0.18, versus chance level = 0.037, *t*_(25)_ = 4.46, *p* < 0.001). However, performance on regular trials was better than on irregular trials (mean proportion difference = 0.73, SD = 0.18, *t*_(25)_ = 20.75, *p* < 0.001). Consistency of mappings across exemplars in the regular set might benefit learning through generalization of mappings across exemplars, while reducing interference between similar exemplars. This was confirmed by comparing behavioural performance in the fourth to the fifth block, when the exemplars were changed. Indeed, performance on regular exemplars showed a consistent increase (proportion correct increase regulars, mean = 0.13, SD = 0.17, *t*_(25)_ = 3.97, *p* < 0.001), which is in line with generalization of acquired knowledge across different exemplars. In contrast, there was no performance increase for irregular trials between block 4 and 5 (proportion correct increase irregulars, mean = 0.01, SD = 0.27, *t*_(25)_ = 0.18, *p* = 0.86) and the performance increase was larger for regular than irregular exemplars (*t*_(25)_ = 2.11, *p* =0.045).

Further evidence of the learning of associative regularity is provided by analysing performance on specific features of the regular objects. By the eight block of acquisition on regular trials, all subjects managed to learn the syllables associated to colour (mean colours correct: 11.85 out of 12, SD = 0.61, chance level = 4, t_(25)_ = 65.30, p<0.001), shape (mean correct: 11.69, SD = 0.84, t_(25)_ = 46.83, p<0.001), and motion (mean correct: 11.38, SD = 0.94, t_(25)_ = 40.00, p<0.001). When probing explicit knowledge of associative regularities at debriefing, all participants could indicate that colour was associated with the first syllable, geometric shape with the second, and movement with the third syllable. Furthermore, when presented with syllables that had occurred in the regular set, along with lures, participants could indicate the meaning of the syllables from the regular set when explicitly probed (mean correct = 11.33 out of 12, SD = 1.52).

### Brain activity associated with feedback-based learning

At the start of learning, participants had no access to prior knowledge of the associative knowledge structure. Therefore, they were required to use information provided during feedback to update their knowledge. To probe brain areas associated with the updating of knowledge, individual learning parameters were fitted to functional MRI data. Learning models were used to capture variations in the shape of individual learning curves. Specifically, the State-Space model provided a better fit to the behavioural data than other learning models (see table 1 and figure 3). The State-Space model was used to analyse the performance pattern across the whole trial history, in order to estimate for each trial the amount that knowledge was updated compared to the next trial. Specifically, a large area (see figure 4, top right panel) spanning the left ventrolateral and lateral prefrontal cortex (*z* = 4.45; cluster size = 239 voxels; MNI coordinates: −38 22 22, p<0.001 FWE corrected for the whole brain) was found to exhibit activity that covaried with the extent to which knowledge is updated based on feedback in the current trial. This region thus may contribute to the updating of representations of the associative knowledge structure. When modeling the accumulated knowledge during feedback, activity was found to covary in several large midline regions, including bilateral medial prefrontal cortex and posterior cingulate cortex, bilateral superior/middle temporal gyrus, and left-lateralized hippocampus, angular gyrus, and somatosensory cortex (see figure 4, lower right panel, and table 2). These regions were more active during feedback presentation when more knowledge had already been accumulated.

**Table 2.** Brain areas where activity significantly correlated trial-by-trial with accumulated knowledge during the feedback trials of the acquisition session. All coordinates are in MNI space.

| Brain region | Cluster size<br>(voxels) | Laterality | Z-score | Local maxima |  |  |
| --- | --- | --- | --- | --- | --- | --- |
|  |  |  |  | x | y | z |
| Medial Prefrontal Cortex | 1740 | L/R | 6.45 | -8 | 55 | 2 |
| Inferior Parietal Cortex | 230 | L | 5.45 | -52 | -62 | 38 |
| Somatosensory Cortex | 260 | R | 5.34 | -65 | -22 | 38 |
| Inferior Parietal Cortex |  |  | 4.89 | 62 | -30 | 45 |
| Inferior Parietal Cortex |  |  | 4.32 | 55 | -32 | 40 |
| Posterior Cingulate Cortex | 853 | L/R | 5.26 | -2 | -12 | 38 |
| Middle Temporal Gyrus | 203 | L | 5.18 | -62 | -18 | -15 |
| Hippocampus | 85 | L | 4.75 | -25 | -18 | -18 |
| Middle/Superior Temporal Gyrus |  | R |  | 60 | -10 | -10 |
|  | 91 |  | 4.66 |  |  |  |

**Figure 4.**
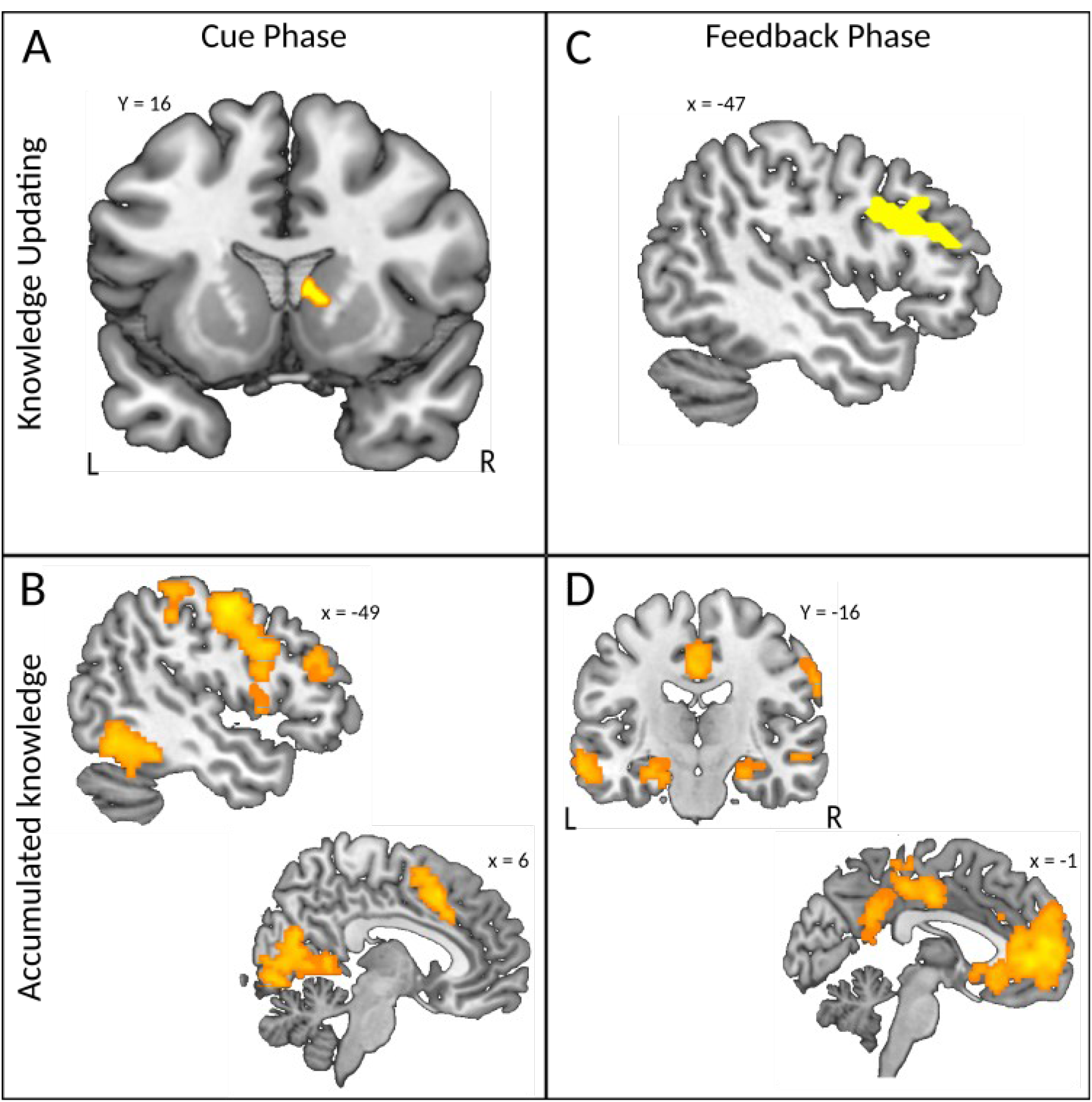
Model-based parametric modulation analysis of learning-related activity. A) Top left panel. Activity in the right caudate nucleus shows a significant correlation with recently updated knowledge available at each test-phase. B) Lower left panel. Regions where activity is correlated with accumulated knowledge at each test-phase, including left inferior frontal gyrus, and visual, motor, and frontoparietal regions. C) Top right panel. Activity in the left ventral and lateral prefrontal cortex is correlated with updated knowledge during the feedback-phase. D) Lower right panel. Regions where activity is correlated with the accumulated knowledge during the feedback-phase, including medial prefrontal cortex, posterior cingulate cortex, left hippocampus, and inferior parietal cortex.

### Brain activity associated with cued retrieval

When presented with the figure during the test-phase, participants were asked to immediately recall the associated tri-syllabic word. Participants updated their knowledge of the associations based on feedback, which they could then apply in the next test-phase. To probe which brain regions aid the deployment of this recently updated knowledge, the updated knowledge parameter was found to track brain activity specifically in the right caudate nucleus (*z* = 4.01; cluster size = 8 voxels; peak MNI coordinates: 8 18 5, p<0.05 corrected for a reduced search volume in an anatomical mask defined by the right caudate nucleus). Thus, the caudate nucleus appears to be involved in the deployment of recently updated knowledge during the test phase (see figure 4, upper left panel). Complementary to this, the knowledge accumulation parameter could be used to probe which regions covary with the total amount of knowledge that had accumulated across previous learning trials and could thus be recruited in the current trial. Several regions were found, including low-level and associative visual regions, primary, premotor and supplementary motor areas, and frontal and parietal cortices (see lower left panel in figure 4, and table 3). Importantly, the left inferior frontal gyrus that we had found to track feedback-based knowledge updating was found to also track accumulated knowledge during the test phase (peak MNI coordinates: −50 12 32, p<0.001 FWE corrected for the whole brain, and peak MNI coordinates: −50 30 22, p<0.001 FWE corrected for the whole brain). In sum, we found a set of regions spanning visual and motor cortex, as well as frontoparietal regions that were more active when more accumulated knowledge was recruited during the test phase.

**Table 3.** Brain areas where activity significantly correlated trial-by-trial with accumulated knowledge during the retrieval trials of the acquisition session. All coordinates are in MNI space.

| Brain region | Cluster size<br>(voxels) | Laterality | Z-score | Local maxima |  |  |
| --- | --- | --- | --- | --- | --- | --- |
|  |  |  |  | x | y | z |
| Premotor Cortex | 460 | L | 5.60 | -48 | -8 | 50 |
| Inferior frontal gyrus |  |  | 5.00 | -50 | 12 | 32 |
| Posterior Cingulate Cortex | 1176 | L/R | 5.46 | 8 | -50 | 5 |
|  |  |  | 5.32 | 2 | -80 | -5 |
| Occipital Cortex |  |  | 4.32 | -8 | -58 | 8 |
| Retrosplenial Cortex |  |  |  |  |  |  |
| Suppl. Motor Area | 473 | L/R | 5.43 | -2 | 10 | 58 |
| Pre-Supp. Motor Area |  |  | 5.14 | 2 | 18 | 45 |
| Anterior Cingulate Cortex |  |  | 4.21 | -8 | 12 | 38 |
| Superior Parietal Cortex | 238 | L | 5.24 | -20 | -62 | 52 |
| Posterior Parietal Cortex |  |  | 4.29 | -18 | -82 | 40 |
| Intraparietal Sulcus |  |  | 3.70 | -28 | -65 | 35 |
| Fusiform Gyrus | 257 | L | 4.89 | -45 | -40 | -20 |
| Occipitotemporal Cortex |  |  | 4.76 | -48 | -60 | -12 |
| Fusiform Gyrus |  |  | 4.54 | -38 | -35 | -25 |
| Occipital Cortex | 73 | L | 4.69 | -35 | -85 | 18 |
| Supplementary Motor Cortex / Broca's Area | 48 | L | 4.44 | -58 | 10 | 0 |
| Thalamus | 44 | L | 4.38 | -10 | -12 | 10 |
| Intraparietal Sulcus | 105 | R | 5.35 | 18 | -65 | 55 |
| Occipitoparietal Cortex |  |  | 3.89 | 18 | -78 | 45 |
| Precuneus |  |  | 3.74 | 20 | -70 | 38 |
| Somatosensory Cortex |  | L | 4.27 | -58 | -18 | 20 |
| Auditory Cortex | 80 |  | 3.57 | -62 | -22 | 10 |
| Inferior Parietal Cortex |  | L | 3.96 | -55 | -30 | 48 |
| Somatosensory Cortex |  |  | 3.91 | -50 | -30 | 58 |
| Intraparietal Sulcus | 52 |  | 3.68 | -45 | -40 | 60 |

### Beta-series extraction

We found regions that responded specifically to the knowledge updating parameters, namely the left inferior frontal gyrus and right caudate nucleus. These regions might thus be involved in the active updating and retrieval of knowledge. Our model detected a wide variety of regions covarying with accumulated knowledge during feedback presentation. Of these regions, the hippocampus is implicated in enabling the generalization across individual learning instances in interaction with the medial prefrontal cortex (Kumaran et al. 2009). It could be that these regions, which covaried with accumulated knowledge during feedback, had some role in generalization of knowledge during feedback-based learning, as more accumulated knowledge might benefit knowledge generalization. To visualize the relative unfolding of activity of these regions across both the test phase and the feedback phase, we extracted the beta time-series from these regions of interest (see figure 5). The left hippocampus and right caudate nucleus were defined by anatomical masks, whereas functional regions found in respective contrast maps (thresholded at p<0.001 uncorrected) were used to define the left inferior frontal gyrus (based on feedback-based knowledge updating) and medial prefrontal cortex (feedback-based accumulated knowledge). The activity across trials estimated by this analysis is displayed in figure 5, and some patterns become apparent upon visual inspection. During feedback-based learning, the left inferior frontal gyrus displays initially high activity, which levels off as more knowledge has already accumulated and less updating takes place. During cue-based retrieval, on the other hand, activity is initially low, but increases as more knowledge accumulates and becomes available to be retrieved. The caudate nucleus displays both initially high activity during feedback-based learning and cue-based retrieval, suggesting it might aid the initial updating and subsequent retrieval of knowledge. Both the hippocampus and medial prefrontal cortex display an increase in activity during feedback-based learning, corroborating the parametric modulation results. During cue-based retrieval, both the hippocampus and the medial prefrontal cortex display an initially high but gradually decreasing activity across retrieval trials as more knowledge is accumulated, suggesting these regions, contrary to expectations, do not become more involved with retrieval as more knowledge is accumulated.

**Figure 5.**
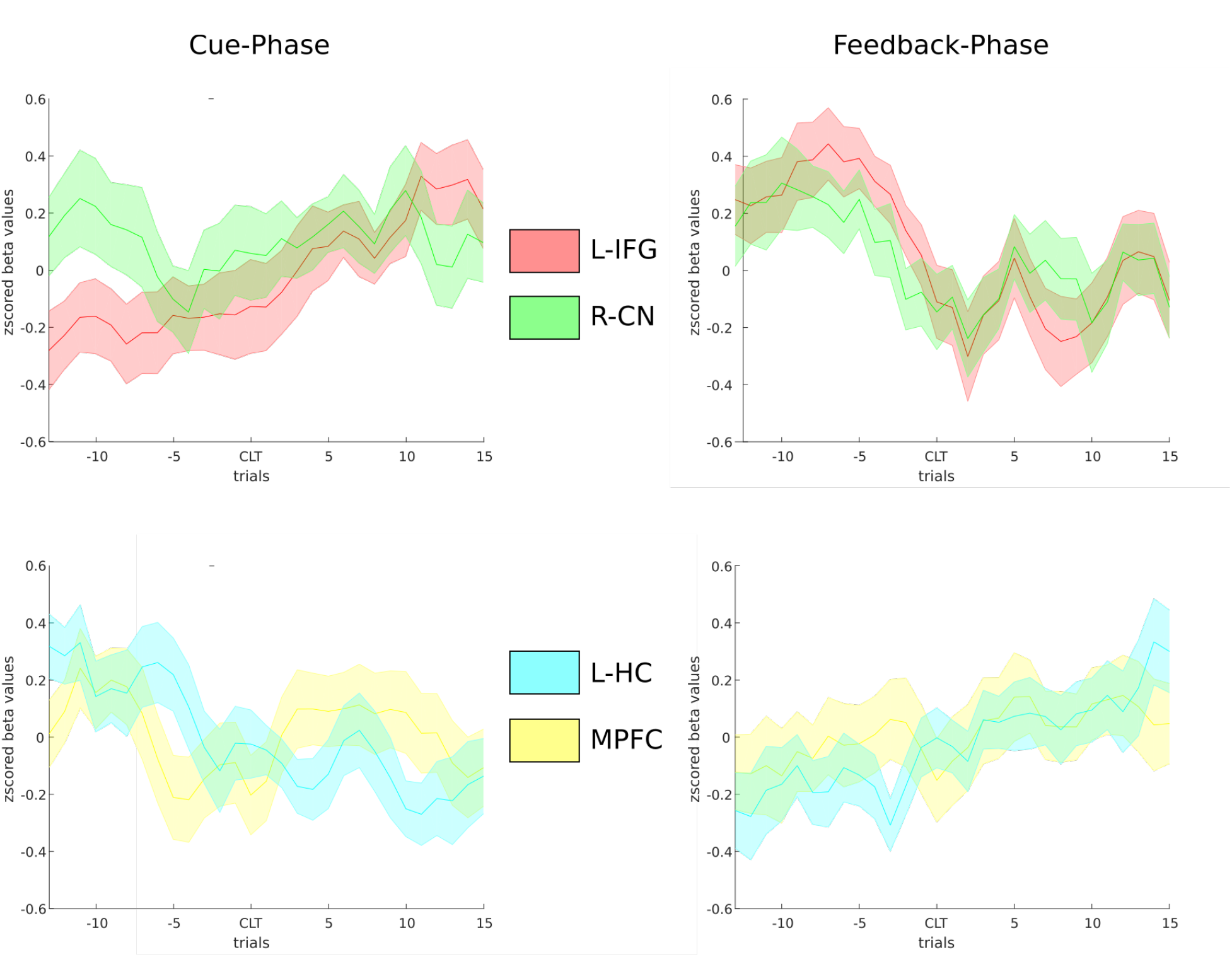
Beta-series analysis of learning. Beta-values were extracted during successive test (left column) respectively successive feedback presentations (right column) across learning. Signal was extracted for the left inferior frontal gyrus (Junctional region that displayed modulation during feedback by knowledge updating; red colours), right caudate nucleus (anatomical mask, this region displayed modulation during the test-phase by recently updated knowledge; green colours) in top row. Signal was also extracted for the left hippocampus (anatomical mask, region displaying modulation during feedback-phase by accumulated knowledge, yellow colours) and medial prefrontal cortex (functional region that displayed modulation during feedback-phase by accumulated knowledge, blue colours. L-IFG = left inferior frontal gyrus, R-CN = right caudate nucleus, L-HC = left hippocampus, MPFC = medial prefrontal cortex.

## Discussion

Few studies have tracked how brain regions dynamically contribute to the gradual acquisition of a complex knowledge structure (Kumaran et al. 2009), and even fewer have disentangled the effects of retrieval and feedback during such learning tasks. The present study addressed this gap by systematically tracking brain activity while learners acquired a novel linguistic associative knowledge structure. The key findings of this study are that feedback-based knowledge updating is associated with activity in the left inferior frontal gyrus. Mirroring this result, the left inferior frontal gyrus displayed a gradual increase in activity during retrieval as more knowledge had accumulated. A complementary signal was found in the right caudate nucleus, where activity during retrieval was found to correlate with the amount of recently updated knowledge that could be retrieved. The results are in line with a model in which the acquisition of linguistic associative knowledge is subserved by the left inferior frontal gyrus, which is initially being supported by fast-learning subcortical regions such as the caudate nucleus. The hippocampus and medial prefrontal cortex, two other regions generally implicated in acquiring generalizable knowledge structures, covaried with accumulated knowledge during feedback presentation. This potentially suggests that as more knowledge accumulates, more generalizable features are extracted through inferential computations performed by the medial prefrontal cortex and hippocampus.

The left inferior frontal gyrus is thus important for linguistic knowledge acquisition. A search on Neurosynth (http://neurosynth.org/) revealed that the peak of the region (x=-38, y=22, y=22) displays the strongest meta-analytical association with the terms “BA44” (z=5.58), “syntax” (z=5.52), “word” (z=4.93) and “semantic” (z=4.82). Indeed, this region has widely been suggested to be involved in language processing (Bookheimer 2002; Friederici 2002; Hagoort 2003; Hagoort et al. 2004). Specifically, it was also implicated in the learning of sequences (Peigneux et al. 1999; Forkstam et al. 2006), and sequences that are hierarchically organized (Gelfand and Bookheimer 2003; Petersson et al. 2004). It has also been argued that the area described here, overlapping with Broca’s area, is involved in retrieving word information from memory and combining them in larger sentence units (Hagoort 2005). The current experiment includes elements from these prior studies, requiring participants to parse hierarchically organized syntactical sequences of feature-syllable associations based on repeated exposures to larger word units (during feedback-based learning), and retrieve word information from memory (during cue-based retrieval).

A particularly interesting finding during retrieval is the initial striatal learning signal, where the right caudate nucleus covaried with the retrieval of recently updated knowledge. This phasic signal during retrieval was complemented by activity in neocortical regions that increased during retrieval as a function of the amount of knowledge that had accumulated. Most of the regions found here were visual, motor, or attentional regions, suggesting they were involved in preparing for the ensuing visuomotor responses and might have been unspecific to the retrieval of knowledge. However, interestingly, the left inferior frontal gyrus, the region that was found to update knowledge on the basis of feedback, was also one of the regions found to display an increase in activity when more accumulated knowledge was available for retrieval, suggesting this region has a function in both active updating of knowledge and subsequent retrieval of accumulated knowledge. This is congruent with a model where this region actively stores the syntactic associative structure: the more this knowledge structure is updated based on feedback, the higher the activity would be in regions that store the knowledge structure, and the more knowledge about the linguistic structure has accumulated across previous trials when recruiting it during retrieval, the more this region would be activated too.

An explanation for a striatal rather than a hippocampal involvement during initial retrieval might be given by the nature of the learning task involved here. The caudate nucleus has generally been found to be important for reward-based reinforcement learning (Haruno et al. 2004; Haruno and Kawato 2006). In a word learning task, signals in the ventral striatum have been related to an implicit or self-generated reward when the meaning of novel word was acquired (Ripollés et al. 2015), while also showing strong connectivity with the more dorsal caudate nucleus and the left inferior frontal gyrus during word learning. In our study, the updating signal observed during the retrieval of knowledge in the caudate nucleus may not have been an explicit reward signal itself (although the feedback provided was probably also experienced as a reward). Rather, the caudate nucleus is more broadly involved in the excitation of correct action schemas and the selection of appropriate sub-goals based on an evaluation of action-outcome contingencies (Grahn et al. 2008). Moreover, prior studies investigating the learning of spatial locations have shown that the hippocampus is involved in incidental configural learning, whereas the dorsal striatum is primarily involved in associative reinforcement learning of stimulus-response contingencies (McDonald and White 1993; Doeller and Burgess 2008; Doeller et al. 2008; Lee et al. 2008). In our task, it might be that participants initially processed associations entirely based on configurations, but then learned to associate certain perceptual features of the figure (color, shape or movement) with a certain syllable at a particular location in the word. This possibility hints at an intriguing trade-off between hippocampal-dependent and striatal-dependent learning, in line with earlier learning studies (using simpler knowledge structures) showing an initial peak in activity in the medial temporal lobe and a somewhat slower build-up of activity in the caudate nucleus (Poldrack et al. 2001). It could be hypothesized that the initial process of discovering regularities requires a recurrent similarity computation by the hippocampus (Kumaran and McClelland 2012). The uncovered component parts of the associative structure are then established in stimulus-response associations between single features and syllables, supported by the striatum. The acquired linguistic associative knowledge structure could then ultimately be stored and retrieved from the left inferior frontal gyrus. Indeed, beta-time series extraction demonstrated an initial increase in retrieval-related activity in both the right caudate nucleus and the hippocampus, as well as the left inferior frontal gyrus, the latter of which continued to increase in activity across retrieval trials (see figure 5).

The hippocampus and medial prefrontal cortex are thought to contribute to the acquisition of generalizable knowledge across trials. Kumaran et al. (2009) found that activity in these regions covaries with accumulated knowledge during learning, and that this learning-related performance was predictive of performance on a transfer test (Kumaran et al. 2009). However, they did not distinguish between retrieval of knowledge when presented with a cue, and the subsequent updating of knowledge based on the provided feedback. In our report, we find that the accumulated knowledge covaries with activity in the hippocampus, medial prefrontal cortex and posterior cingulate cortex, similar to the report by Kumaran et al. However, we find this relation specifically during the presentation of feedback. Even though the associative structure acquired in our task consisted of linguistic material and is more complex, the basic computations needed to acquire the associative knowledge structure are quite similar (Kumaran and McClelland 2012). Thus, despite the fact that recurrent similarity computations subserved by the hippocampus and its connectivity with the medial prefrontal cortex might initially already take place to enable an acceleration of learning, these computations are continuously needed to acquire all associative rules (associations between single feature and syllables, and the larger hierarchical associative structure) until an optimal performance is reached. Thus, activity in the medial prefrontal cortex and the hippocampus would track recurrent similarity computations underlying generalization of knowledge, which would track accumulated knowledge during feedback presentation. Alternative factors might be postulated to explain a relation of these brain regions and accumulated knowledge, such as reward processing: the accumulated knowledge parameter scales with the correctness of feedback, potentially eliciting an implicit reward signal. Moreover, various other regions in posterior cingulate cortex, temporal and parietal cortices were also found to covary with accumulated knowledge. More detailed behavioural paradigms are needed that can be performed in short learning sessions to better assess the neural dynamics of these learning systems.

A cautionary note should be made. It could be argued that the learning benefit for regular trials is not solely due to an ability to generalize knowledge across trials. For instance, in irregular trials, feature-syllable associations are constantly changing, potentially creating interference between associations that partially share the same features. However, increased interference across trials is an inherent outcome of reduced consistency. Similarly, consistency across trials reduces interference, and therefore inherently promotes generalization across trials. Furthermore, not only did we establish that generalization took place by comparing learning performance in regular and irregular trials, we also find that participants generalized between the first half and the second half of the learning session when exemplars were switched.

In conclusion, this study provides novel insights into how linguistic associative knowledge is acquired by systematically tracking schematic knowledge formation while participants were learning an abstract artificial language organized by higher-order associative regularity. During learning, we find signals in the left inferior frontal gyrus, which responds both to feedback-based knowledge updating, and available accumulated knowledge during retrieval. A complementary signal is found in the caudate nucleus, where activity correlates with the availability of recently acquired knowledge during retrieval, suggesting it initially supports the retrieval of knowledge. Furthermore, we find that activity in a set of regions, including the medial prefrontal cortex and hippocampus, scaled with accumulated knowledge during feedback presentation, which might be indicative of increased generalization of features of the hierarchical knowledge structure. Together, these results provide a mechanistic insight into how linguistic associative knowledge is acquired by generalization across repeated learning experiences.

**Author contributions:** R.M.WJ.B., M.vdL., M.T.R.vK., R.G.M.M. & G.F. designed research; R.B. performed research; D.N. & J.M. contributed analytic tools; R.B.,D.N. & J.M. analyzed data; R.M.WJ.B. wrote the manuscript; all authors commented on and provided feedback on drafts of the paper.

## Acknowledgement

R.M.W.J.B., M.v.d.L., R.G.M.M. and G.F are supported by a grant from the European Research Council (“Neuroschema” ERC R0001075). G.F., D.A.N. & J.M. are supported by a grant from the NWO (“Language in Interaction”). We thank Isabella Wagner, Peter Koopmans and Tessa van Leeuwen for help with programming task scripts and Arjen Stolk and Nils Miller for inspiring discussions. M.T.R.vK. is now at Faculty of Behavioural and Movement Sciences, Institute for Brain and Behaviour, Vrije Universiteit Amsterdam, Amsterdam, the Netherlands

## References

Arppe A, Hendrix P, Milin P, Baayen RH, Sering T, Shaoul C. 2015. ndl: Naive Discriminative Learning. R package version 0.2.17.

Barsalou LW. 2009. Simulation, situated conceptualization, and prediction. Philos Trans R Soc London B Biol Sci. 364:1281–1289.

Bartlett FC. 1932. Remembering: An experimental and social study, Cambridge: Cambridge University.

Bookheimer S. 2002. Functional MRI of language: new approaches to understanding the cortical organization of semantic processing. Annu Rev Neurosci. 25:151–188.

Brady TF, Oliva A. 2008. Statistical Learning Using Real-World Scenes: Extracting Categorical Regularities Without Conscious Intent. Psychol Sci. 19:678–685.

Brainard DH. 1997. The Psychophysics Toolbox. Spat Vis. 10:433–436.

Chessa AG, Murre JMJ. 2007. A Neurocognitive Model of Advertisement Content and Brand Name Recall. Mark Sci. 26:130–141.

Dalton MA, Weickert TW, Hodges JR, Piguet O, Hornberger M. 2013. Impaired acquisition rates of probabilistic associative learning in frontotemporal dementia is associated with fronto-striatal atrophy. NeuroImage Clin. 2:56–62.

Danks D. 2003. Equilibria of the Rescorla-Wagner model. J Math Psychol. 47:109–121.

Daw ND, Niv Y, Dayan P. 2005. Uncertainty-based competition between prefrontal and dorsolateral striatal systems for behavioral control. Nat Neurosci. 8:1704–1711.

Doeller CF, Burgess N. 2008. Distinct error-correcting and incidental learning of location relative to landmarks and boundaries. Proc Natl Acad Sci U S A. 105:5909–5914.

Doeller CF, King JA, Burgess N. 2008. Parallel striatal and hippocampal systems for landmarks and boundaries in spatial memory. Proc Natl Acad Sci. 105:5915–5920.

Forkstam C, Hagoort P, Fernández G, Ingvar M, Petersson KM. 2006. Neural correlates of artificial syntactic structure classification. Neuroimage. 32:956–967.

Friederici AD. 2002. Towards a neural basis of auditory sentence processing. Trends Cogn Sci. 6:7884.

Gelfand JR, Bookheimer SY. 2003. Dissociating neural mechanisms of temporal sequencing and processing phonemes. Neuron. 38:831–842.

Ghosh VE, Gilboa A. 2014. What is a memory schema? A historical perspective on current neuroscience literature. Neuropsychologia. 53:104–114.

Grahn JA, Parkinson JA, Owen AM. 2008. The cognitive functions of the caudate nucleus. Prog Neurobiol. 86:141–155.

Hagoort P. 2003. How the brain solves the binding problem for language: a neurocomputational model of syntactic processing. Neuroimage. 20:S18–S29.

Hagoort P. 2005. On Broca, brain, and binding: a new framework. Trends Cogn Sci. 9:416–423.

Hagoort P, Hald L, Bastiaansen M, Petersson KM. 2004. Integration of word meaning and world knowledge in language comprehension. Science (80- ). 304:438–441.

Halai AD, Parkes LM, Welbourne SR. 2015. Dual-echo fMRI can detect activations in inferior temporal lobe during intelligible speech comprehension. Neuroimage. 122:214–221.

Haruno M, Kawato M. 2006. Different neural correlates of reward expectation and reward expectation error in the putamen and caudate nucleus during stimulus-action-reward association learning. J Neurophysiol. 95:948–959.

Haruno M, Kuroda T, Doya K, Toyama K, Kimura M, Samejima K, Imamizu H, Kawato M. 2004. A neural correlate of reward-based behavioral learning in caudate nucleus: a functional magnetic resonance imaging study of a stochastic decision task. J Neurosci. 24:1660–1665.

Holl AK, Wilkinson L, Tabrizi SJ, Painold A, Jahanshahi M. 2012. Probabilistic classification learning with corrective feedback is selectively impaired in early Huntington’s disease–Evidence for the role of the striatum in learning with feedback. Neuropsychologia. 50:2176–2186.

Hyndman R, Athanasopoulos G, Razbash S, Schmidt D, Zhou Z, Khan Y. 2013. forecast: Forecasting functions for time series and linear models [R package version 4.03].

Kirby S, Cornish H, Smith K. 2008. Cumulative cultural evolution in the laboratory: an experimental approach to the origins of structure in human language. Proc Natl Acad Sci U S A. 105:10681–10686.

Knowlton BJ, Squire LR, Gluck MA. 1994. Probabilistic classification learning in amnesia. Learn Mem. 1:106–120.

Kumaran D, McClelland JL. 2012. Generalization through the recurrent interaction of episodic memories: a model of the hippocampal system. Psychol Rev. 119:573–616.

Kumaran D, Summerfield JJ, Hassabis D, Maguire EA. 2009. Tracking the emergence of conceptual knowledge during human decision making. Neuron. 63:889–901.

Lee AS, Duman RS, Pittenger C. 2008. A double dissociation revealing bidirectional competition between striatum and hippocampus during learning. Proc Natl Acad Sci U S A. 105:17163–17168.

Markman EM, Hutchinson JE. 1984. Children’s sensitivity to constraints on word meaning: Taxonomic versus thematic relations. Cogn Psychol. 16:1–27.

McClelland JL, McNaughton BL, O’Reilly RC. 1995. Why there are complementary learning systems in the hippocampus and neocortex: insights from the successes and failures of connectionist models of learning and memory. Psychol Rev. 102:419–457.

McDonald RJ, White NM. 1993. A triple dissociation of memory systems: hippocampus, amygdala, and dorsal striatum. Behav Neurosci. 107:3–22.

Miller EK, Freedman DJ, Wallis JD. 2002. The prefrontal cortex: categories, concepts and cognition. Philos Trans R Soc London B Biol Sci. 357:1123–1136.

Murre JMJ. 2013. S-shaped learning curves. Psychon Bull Rev. 21:344–356.

O’Reilly RC, Norman KA. 2002. Hippocampal and neocortical contributions to memory: advances in the complementary learning systems framework. Trends Cogn Sci. 6:505–510.

Pasupathy A, Miller EK. 2005. Different time courses of learning-related activity in the prefrontal cortex and striatum. Nature. 433:873–876.

Peigneux P, Maquet P, Van der Linden M, Meulemans T, Degueldre C, Delfiore G, Luxen A, Cleeremans A, Franck G. 1999. Left inferior frontal cortex is involved in probabilistic serial reaction time learning. Brain Cogn. 40:215–219.

Petersson KM, Forkstam C, Ingvar M. 2004. Artificial syntactic violations activate Broca’s region. Cogn Sci. 28:383–407.

Poldrack RA, Clark J, Pare-Blagoev EJ, Shohamy D, Creso Moyano J, Myers C, Gluck MA. 2001. Interactive memory systems in the human brain. Nature. 414:546–550.

Rescorla RA, Wagner AR. 1972. A theory of Pavlovian conditioning: Variations in the effectiveness of reinforcement and nonreinforcement. In: Classical conditioning: Current research and theory.

Ripollés P, Marco-Pallarés J, Hielscher U, Mestres-Missé A, Tempelmann C, Heinze H-J, Rodríguez-Fornells A, Noesselt T. 2015. The Role of Reward in Word Learning and Its Implications for Language Acquisition. Curr Biol. 24:2606–2611.

Schwarz G. 1978. Estimating the dimension of a model. Ann Stat. 6:461–464.

Seger CA, Cincotta CM. 2006. Dynamics of Frontal, Striatal, and Hippocampal Systems during Rule Learning. Cereb Cortex. 16:1546–1555.

Seger CA, Miller EK. 2010. Category learning in the brain. Annu Rev Neurosci. 33:203–219.

Seger CA, Poldrack RA, Prabhakaran V, Zhao M, Glover GH, Gabrieli JDE. 2000. Hemispheric asymmetries and individual differences in visual concept learning as measured by functional MRI. Neuropsychologia. 38:1316–1324.

Smith AC, Frank LM, Wirth S, Yanike M, Hu D, Kubota Y, Graybiel AM, Suzuki WA, Brown EN. 2004. Dynamic Analysis of Learning in Behavioral Experiments. J Neurosci. 24:447–461.

Tse D, Langston RF, Kakeyama M, Bethus I, Spooner PA, Wood ER, Witter MP, Morris RGM. 2007. Schemas and Memory Consolidation. Sci. 316:76–82.

Tse D, Takeuchi T, Kakeyama M, Kajii Y, Okuno H, Tohyama C, Bito H, Morris RGM. 2011. Schema-Dependent Gene Activation and Memory Encoding in Neocortex. Sci. 333:891–895.

van der Linden M, Berkers RMWJ, Morris RGM, Fernández G. 2017. Angular gyrus involvement at encoding and retrieval is associated with durable but less specific memories. J Neurosci. 37.

van Kesteren MTR, Fernández G, Norris DG, Hermans EJ. 2010. Persistent schema-dependent hippocampal-neocortical connectivity during memory encoding and postencoding rest in humans. ProcNatl Acad Sci. 107:7550–7555.

van Kesteren MTR, Rijpkema M, Ruiter DJ, Morris RGM, Fernández G. 2014. Building on prior knowledge: schema-dependent encoding processes relate to academic performance. J Cogn Neurosci. 26:2250–2261.

van Kesteren MTR, Ruiter DJ, Fernández G, Henson RN. 2012. How schema and novelty augment memory formation. Trends Neurosci. 35:211–219.

Wagner IC, van Buuren M, Kroes MCW, Gutteling TP, van der Linden M, Morris RG, Fernández G. 2015. Schematic memory components converge within angular gyrus during retrieval. Elife. 4.

